# Pax2a regulates angiogenesis to facilitate mmp2-dependent basement membrane remodeling of the optic fissure

**DOI:** 10.1101/678185

**Authors:** Megan L. Weaver, W. P Piedade, N.N Meshram, J.K. Famulski

## Abstract

Vertebrate retinal development requires timely and precise fusion of the optic fissure (OF). Recent studies have suggested hyaloid vasculature to be involved in optic fissure fusion. In order to examine this link, we analyzed OF fusion and hyaloid vasculogenesis in the zebrafish pax2a^noi^ mutant line. We determined that OF basement membrane (BM) remodeling, normally preceded by F-actin accumulation is mis-regulated in pax2a^−/−^ embryos. Comparing transcriptomic profiles of pax2a^−/−^ and wildtype eyes we discovered a novel connection between regulation of angiogenesis and fusion. Pax2a^−/−^ eyes exhibited a significant reduction of *talin1* expression, a regulator of hyaloid vasculature, in addition to increased expression of an anti-angiogenic protease, *adamts1*. Using 3D and live imaging we observed reduced OF hyaloid vascularization in pax2a^−/−^ embryos. Additionally, pharmacological inhibition of VEGF signaling or *adamts1* mRNA overexpression phenocopied the pax2a^−/−^ vasculature, F-actin and BM remodeling phenotypes. Finally, we show that hyaloid vasculature expresses *mmp2* which is necessary for remodeling the fissure BM. Taken together we propose a pax2a driven mechanism that restricts anti-angiogenic activity of adamts1 enabling hyaloid vasculature invasion of the OF and delivery of the BM remodeler mmp2.

## INTRODUCTION

Ocular development is a highly conserved process amongst vertebrate species. Assembly of the hemispherical, retinal structure from an initially flat sheet of cells requires many complex morphogenetic movements. One such morphogenetic movement involves the invagination of the optic vesicle which results in a fissure forming at the ventral region of the developing retina. This fissure, known as the choroid or optic fissure (OF), enables hyaloid vasculature cell migration into the developing retina and subsequent establishment of the hyaloid vasculature. Hyaloid vasculature is a temporary circulatory system required for ocular development, and in most cases will degenerate once mature blood vessels begin to grow (1–4). As soon as the hyaloid vasculature has been established, the two opposing retinal epithelial sheets of the optic fissure will undergo fusion. Thereby, they encase the ganglion cell axons localized in the optic stalk and complete retinal morphogenesis. Failure of optic fissure fusion leads to a congenital blinding disorder known as coloboma (5–7). Coloboma is a prevalent cause of pediatric blindness, accounting for approximately 10% of cases worldwide (6, 8). This makes it one of the leading causes of pediatric blindness. Coloboma is a spectrum disorder presenting unilaterally or bilaterally and ranging in severity from minor visual impairment, to complete blindness in the affected eye (9). This spectrum of severity is associated with the location and degree to which the choroid fissure was able to fuse and the severity of subsequent loss of ganglion cell axons (7).

Coloboma has been studied for many decades in many different species. Work over this time has led to a general outline of the signaling and morphogenetic pathways required for proper optic fissure formation and fusion (recently reviewed in (10)). In particular, opposing action of bone morphogenetic protein (BMP) and sonic hedgehog (Shh) signaling establishing the dorsal-ventral pattern of the optic vesicle and ensuring proper expression of optic stalk and OF regulators pax2, vax1 and vax2 (11, 12). However, the actual molecular mechanisms driving optic fissure fusion remains largely unknown. The process of epithelial tissue fusion is not unique to the eye and occurs throughout development, encompassing neural tube closure, palatal shelf formation and eyelid development (13). Epithelial fusion has been studied for over a century, and is known to involve transcriptional regulation, cell signaling pathways and morphogen gradients (14, 15). The actin cytoskeleton is thought to be a crucial component of the machinery driving epithelial fusion in many tissues (16). The importance of the actin cytoskeleton during epithelial fusion involves lamellipodial and filopodial projections between the two opposing epithelia. These help to “zipper” the cells together to form a single continuous sheet. When lamellipodial and filopodial projections are precluded, epithelial fusion often fails (17). Lamellipodia and filopodia have been observed during OF fusion almost 3 decades ago (18–20). However, the functional and regulatory mechanisms behind these projections remain unknown. Another cellular mechanism known to be directly involved in epithelial fusion is the remodeling of the basement membrane (BM). During epithelial fusion, the BM acts as a physical barrier restricting the establishment of cell-cell contacts, which must be removed in order to complete fusion. Recent work from several labs, working on different species, has characterized progressive removal of the BM during OF fusion (21–24). However, the molecular mechanisms facilitating this process, in particular BM remodeling, also remain largely unknown. It was recently suggested that hyaloid vasculature cells migrating through the OF could potentially signal or facilitate BM remodeling (22). Hyaloid vasculature nourishes the developing retina and lens while connecting to the choroid vasculature for proper blood flow (3). Hyaloid vasculogenesis takes advantage of an open optic fissure so that vasculature cells can migrate into the developing optic cup. Once optic fissure fusion is completed, hyaloid vasculature is fully established. James *et al.* 2016, showed that mutations in zebrafish *talin1*, an actin cytoskeleton scaffolding protein known to be required for endothelial cell migration (25), result in OF fusion defects. Theirs, and previous studies also indicated that *cloche* mutants, which lack all early hyaloid vasculature, have delayed basement membrane breakdown in the region of the choroid fissure (22, 26). Since the hyaloid vasculature requires an open fissure to complete establishment of its network, it has been proposed that migrating hyaloid vasculature cells may regulate the timing of fissure fusion. This mechanism could potentially involve vasculature-mediated activation of the fusion machinery within the retinal rim cells, or direct supply of molecular factors, such as matrix proteases (27). In additional support of this hypothesis, recent optic cup transplantation experiments in zebrafish embryos confirm that in the absence of hyaloid vasculature, ectopic retinal OFs fail to initiate fusion (28). Hence, there is a clear link between OF fusion and hyaloid vasculature.

In our current study, we have undertaken a detailed analysis of zebrafish OF fusion comparing WT and the pax2a^−/−^ coloboma model (29). In particular, we have characterized the timing of BM remodeling, cytoskeletal responses, morphological apposition and hyaloid vascularization. Using transcriptomic analysis to compare WT and pax2a^−/−^ eyes we discovered mis-regulation in angiogenic signaling, resulting in decreased OF hyaloid vascularization and OF fusion failure. In particular we found that pax2a^−/−^ embryos exhibit a decrease in *talin1* expression coupled with increased expression of an anti-angiogenic protease, *a disintegrin and metalloproteinase with thrombospondin motifs 1 (adamts1*), (30, 31). Modulation of vascularization via pharmacological inhibition of VEGF signaling or overexpression of adamts1 both phenocopied the pax2a^−/−^ hyaloid vasculature and coloboma phenotypes. Mechanistically, we also show that hyaloid vasculature is a source of mmp2, mmp14a and mmp14b and that mmp2 activity is necessary for OF BM remodeling. Taken together, we propose a novel pathway for the regulation of OF fusion where pax2a mediates proper expression of *adamts1* in order to facilitate hyaloid vasculature recruitment to the OF and subsequent vasculature supplied mmp2-dependent BM remodeling.

## RESULTS

### Optic fissure basement membrane remodeling is preceded by F-actin accumulation

Several recent studies have undertaken a detailed time course to map out the exact timing of optic fissure fusion in numerous species, including zebrafish (21–24). Overall, in zebrafish, the data point to ∼32-36hpf as the time of OF fusion initiation, as observed by BM remodeling. To decipher the molecular mechanisms regulating OF fusion we also performed a detailed time course analysis of OF BM remodeling using laminin immunohistochemistry (IHC) and additionally analyzed F-actin. Our goal was to determine whether changes in F-actin levels in the OF were correlative with fusion. James *et al*, 2016 had recently suggested that infiltrating vasculature endothelial cells migrating through the fissure could be a source of signal for fissure fusion. In fact, they have shown that interactions between vasculature and the fissure resulted in an increase of F-actin (22). As such, we performed whole mount IHC for laminin deposition in embryos starting at 24hpf. We sampled every 4 hours up to 48hpf and then at 56 and 72hpf (Fig 1A). To analyze the progression of the fusion process, we quantified the laminin signal within the OF from 3D confocal scans. Regions of the BM outside the OF and juxtaposed to the developing lens were used to normalize the laminin signal (Fig S1A). We quantified distal, medial and proximal regions of the fissure (Fig S1B,C). When comparing results from the three regions we did not observe significant differences in the distribution of laminin or F-actin. Going forward we focused on quantifying laminin signal in the central proximal region (medial) of the fissure (Fig S1B). This region has been previously shown to represent the site of fusion initiation and is easily identified in 24-72hpf embryos. In agreement with recent studies in zebrafish, we found that OF fusion initiates at ∼32-36hpf and does so in a central region of the fissure, subsequently proceeding proximally and distally (Fig 1B). By 48hpf, we observed most of the laminin signal was removed from the OF, and by 72hpf, little to no laminin persists in the region (Fig 1A, B).

**Figure 1:**
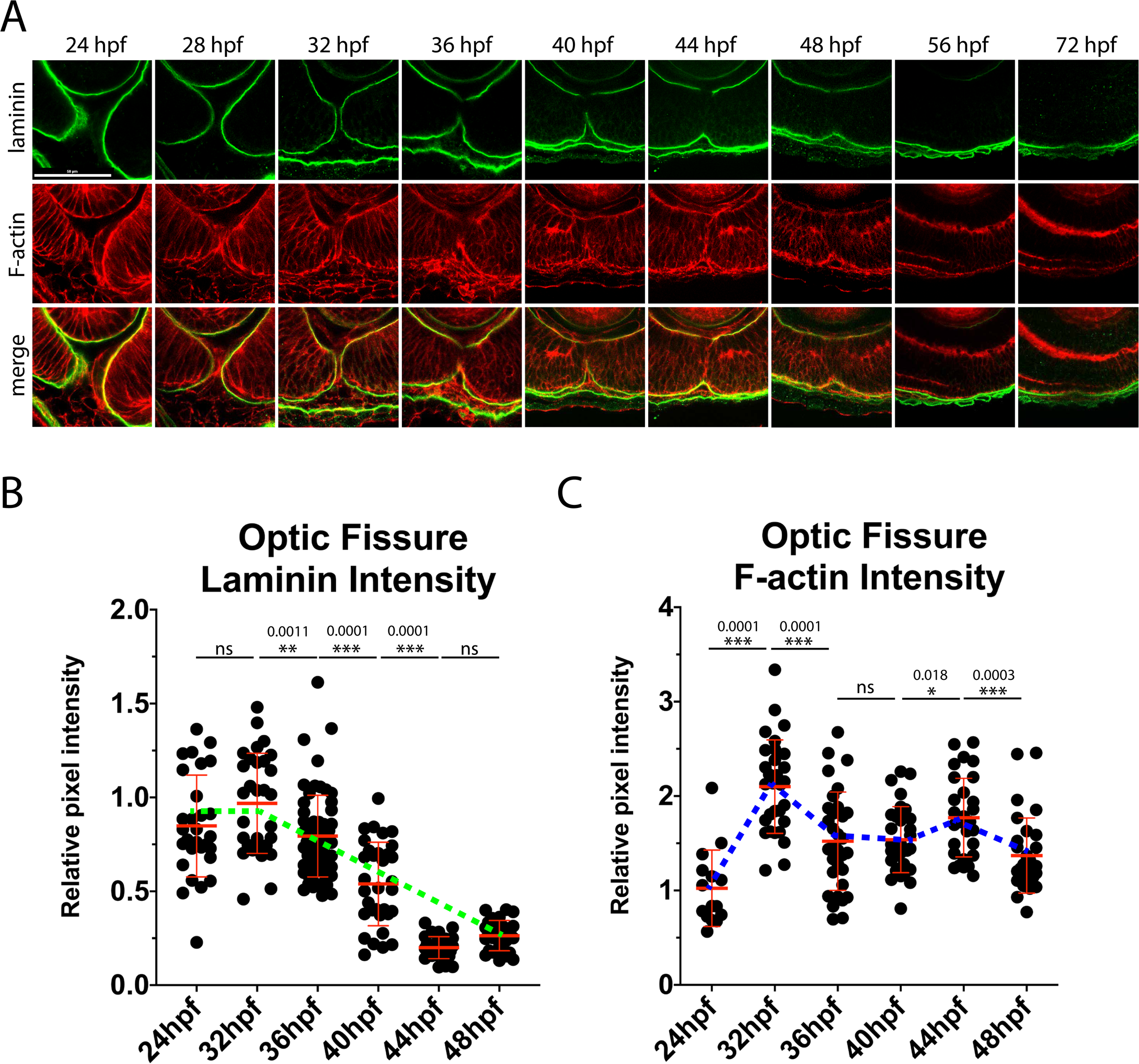
An increase in F-actin dynamics precedes laminin remodeling during optic fissure fusion. **A)** Whole mount immunohistochemistry was used to simultaneously visualize F-actin (red) and laminin (green) during optic fissure fusion, 24-72hpf. Central-proximal sections obtained using confocal imaging were collected and quantified. Scale bar = 50um. **B)** Quantification of laminin signal intensity within the optic fissure, normalized to regions of laminin staining juxtaposed to the lens. Relative pixel intensities are displayed. The green dotted line depicts the trend in laminin intensity over time. ANOVA p<0.0001. **C)** Quantification of F-actin intensity (phalloidin staining) within the optic fissure, normalized to regions of F-actin signal within the lobe of the retina. Relative pixel intensities are displayed. The blue dotted line represents the trend in F-actin intensity over time. ANOVA p<0.0001.

To track F-actin levels during fusion we stained the laminin labeled embryos with phalloidin and quantified the signal within the fissure. To normalize phalloidin staining intensity we used an interior region of the retina (Fig S1A). Our analysis revealed an increase in OF associated F-actin signal between 24-32hpf, preceding the time that BM remodeling is initiated (Fig 1A,B). We also noticed a second increase in F-actin around 44hpf, a time of active fusion (Fig 1C). The observed increase of F-actin signal correlates with a) the timing of hyaloid vasculature migration (24-32hpf) into the fissure and b) the expected timing of actin protrusions involved in epithelial fusion (36-48hpf). When measuring distance between the fusing lobes of the retina we observed apposition to complete around 36-40hpf (Fig S2). Taken together, we conclude that both apposition and BM remodeling are preceded by an increase of F-actin within the OF.

### Optic fissure fusion mechanics are disrupted upon loss of pax2a function

In order to test our theory that an increase in F-actin is involved in the initiation of OF fusion we compared our findings to an established model of coloboma, the pax2a^noi^ line (29). Pax2 is required for OF fusion in several model systems and has been documented in human coloboma cases (32–34). The noi mutation is predicted to result in a loss of pax2a function due to a premature stop codon at position 198 (35). Originally characterized for their no-isthmus phenotype, pax2a^noi/noi^ embryos (referred to as pax2a^−/−^ from this point on) elicit a fully penetrant unfused OF. Unfortunately, homozygous mutants are not viable and do not enable the study of coloboma at juvenile or adult stages. We hypothesized that examining the molecular events leading to OF fusion in the pax2a^noi^ system would inform us whether these events are functionally important. Similar to our WT study, we examined apposition, laminin and F-actin levels throughout the time of OF fusion (Fig 2A, Fig S1C). In stark contrast to WT, pax2a^−/−^ embryos did not exhibit a significant decrease in laminin signal between 32-48hpf. In fact, as expected, laminin appears to be largely retained in the OF of pax2a^−/−^ embryos up to 72hpf (Fig 2B). Previous studies in pax2a null mice had indicated failure of BM remodeling and retention of laminin (34, 36, 37). While there is a moderate decrease in laminin signal in pax2a^−/−^ embryos from 32-36hpf, this result is significantly different from WT. IHC images clearly show that laminin persists in the fissure up to and including 72hpf (Fig 2A). When examining OF F-actin levels in pax2a^−/−^ embryos we did not detect the expected increase between 24 and 32hpf (Fig 2C). We observed F-actin levels fluctuating between 36 and 44hpf and finally dropping to WT amounts by 48hpf. Similar results were also observed when quantifying distal and proximal regions of the fissure (Fig S1D). Our data therefore suggest that the absence of OF F-actin accumulation between 24 and 32hpf, correlates with failure to initiate fusion. Lastly, when examining retinal lobe distance in pax2a^−/−^ embryos we did not observe any significant defects in the timing of apposition (Fig S2). Taken together, we conclude that in the absence of pax2a F-actin fails to accumulate in the OF and this correlates with the failure to remodel the BM. Based on our analysis of apposition, we also conclude that pax2a does not regulate morphogenetic movements of the retinal lobes. In summary, we propose that accumulation of F-actin is necessary for the initiation of OF BM remodeling. We therefore next sought to investigate the mechanism of OF F-actin accumulation.

**Figure 2:**
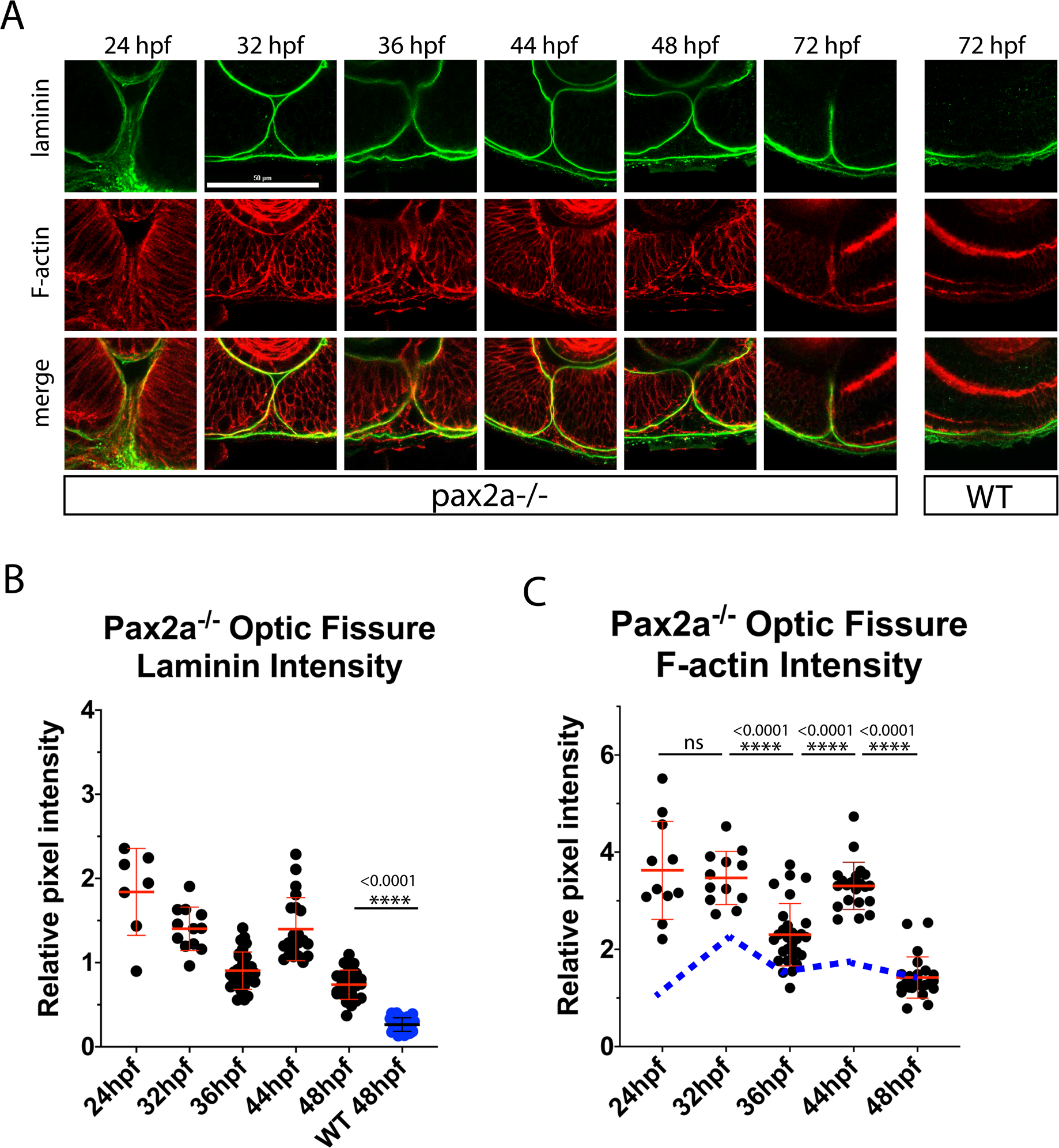
Pax2 mutant embryos lack F-actin activity increase and laminin remodeling within the optic fissure. **A)** Whole mount Immunohistochemistry was used to simultaneously visualize F-actin (red) and laminin (green) during optic fissure fusion, 24-72hpf, in pax2^−/−^ embryos. Central-proximal regions of the optic fissure are displayed. Scale bar = 50um **B)** Quantification of laminin signal intensity within the central-proximal region of the optic fissure, normalized to regions of laminin staining juxtaposed to the lens. Relative pixel intensities are displayed. ANOVA p<0.0001. **C)** Quantification of F-actin intensity (phalloidin staining) within the central-proximal region of the optic fissure, normalized to regions of F-actin signal within the lobe of the retina. Relative pixel intensities are displayed. The blue dotted line represents the trend of F-actin intensity observed in WT embryos. ANOVA p<0.0001.

### Pax2a retinal transcriptomic profile during optic fissure fusion

Although pax2 has been studied in many systems, there are currently no obvious transcriptional targets of pax2 that would directly point to mechanistic regulation of OF fusion. In an effort to understand the absence of OF BM remodeling and mis-regulation of F-actin observed in pax2a^−/−^ embryos we compared whole eye transcriptomic profiles between WT and pax2a^−/−^ embryos. Eyes from 48hpf WT and pax2a^−/−^ embryos were isolated and total RNA was subsequently purified and sequenced using Illumina sequencing. 48hpf was chosen for our experimental time point as pax2a^−/−^ embryos are easily phenotyped at this age thanks to observable heart malformations. When comparing three replicates for WT and pax2a^−/−^ embryos using RSEM software, we detected 1215 transcripts significantly upregulated (>95% confidence interval) and 1202 transcripts significantly downregulated (Fig 3A and 3D). Gene ontology analysis indicated a wide spread of biological function being affected, including cytoskeletal signaling, adhesion and developmental processes (Fig 3B, E). In fact, the distribution of biological function between up and downregulated genes was highly similar (Fig 3B, D). A list of the top 20 up and down regulated genes is outlined in Figure 3C and F. Results of all the statistically significant up and downregulated genes identified in our assay are presented as supplementary data (Table 1 and Table 2).

**Figure 3:**
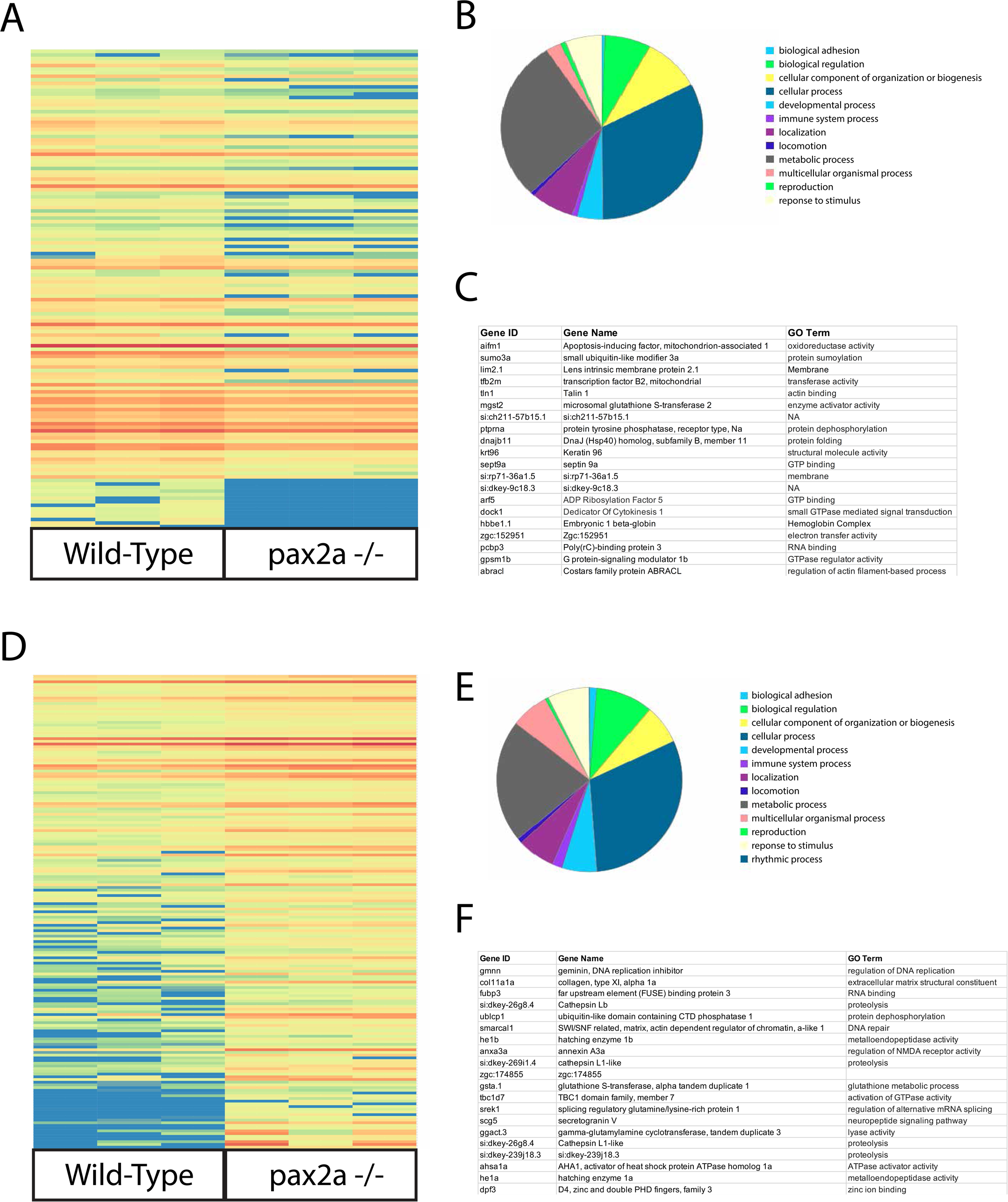
Retinal transcriptomic comparison between WT and pax2a^−/−^. **A)** Heat map representing transcripts found to be significantly down regulated in pax2a^−/−^ embryos. **B)** Gene ontology for pax2a^−/−^ downregulated transcripts. **C)** Top 20 downregulated transcripts are outlined in order of fold change. **D)** Heat map representing transcripts found to be significantly upregulated in pax2a^−/−^ embryos. **E)** Gene ontology for pax2a^−/−^ upregulated transcripts. **F)** Top 20 upregulated transcripts are outlined in order of fold change.

When examining our list of downregulated targets, one, *talin1* (tln1), stood out in particular. Not only was *tln1* one of the most downregulated targets (∼200 fold decrease) it was also recently shown to be necessary for the migration of hyaloid vasculature cells into the OF and subsequent fusion (22). As such, we next sought to investigate the relationship between pax2a, tln1, vasculogenesis and optic fissure fusion mechanics.

### Loss of pax2a function leads to reduced hyaloid vasculature within the optic fissure

To validate our RNAseq data, we performed wholemount *in situ* hybridization (WISH) for *tln1* comparing WT siblings to pax2a^−/−^ mutant embryos. WT expression of *tln1* was observed in the optic fissure between 28 and 48hpf coinciding with pax2a expression (Fig 4A, Fig S3). In pax2a^−/−^ embryos, OF *tln1* expression appears significantly reduced compared to WT while retaining similar expression in periocular regions (Fig 4A). *Tln1* expression is also reduced in the mid brain-hind brain boundary, another region of strong *pax2a* expression (Fig 4A, S3A). Our WISH data was further supported by qPCR results showing a significant decrase in *tln1* expression at 32hpf (Fig 4B). To determine whether the reduction of *tln1* expression in pax2a^−/−^ embryos had functional consequences on hyaloid vasculogenesis we examined pax2a^−/−^ Tg[kdrl:mCherry] embryos. Using 3D *in vivo* time-lapse confocal microscopy we recorded migration of mCherry expressing cells through the OF from 24 to 30hpf (Fig 4B, Movie 1). At 24hpf, both WT and pax2a^−/−^ embryos contain mCherry expressing cells within the fissure. However, over the next six hours of imaging it is apparent that pax2a^−/−^ embryos have significantly fewer mCherry expressing cells pass through the OF (Fig 4C, Movie 1). To quantify this effect, we fixed WT and pax2a^−/−^ Tg[kdrl:mCherry] embryos at 24, 32, 36 and 48hpf, collected 3D confocal stacks and counted the number of mCherry positive cells found within the OF (Fig 4D). The data indicate that in pax2a^−/−^ embryos there is a significant reduction in the number of mCherry expressing cells at both 32 and 36hpf (Fig 4E). Furthermore, using 3D rendering, we noted that in 48hpf pax2a^−/−^ embryos the hyaloid vasculature established in the back of the lens is reduced in size and lacks proper connections to the newly forming choroidal and superficial vasculature systems (Movie 2-3). Overall, our data indicate that pax2a^−/−^ embryos exhibit decreased expression of *tln1* and impaired hyaloid vascularization of the optic fissure.

**Figure 4:**
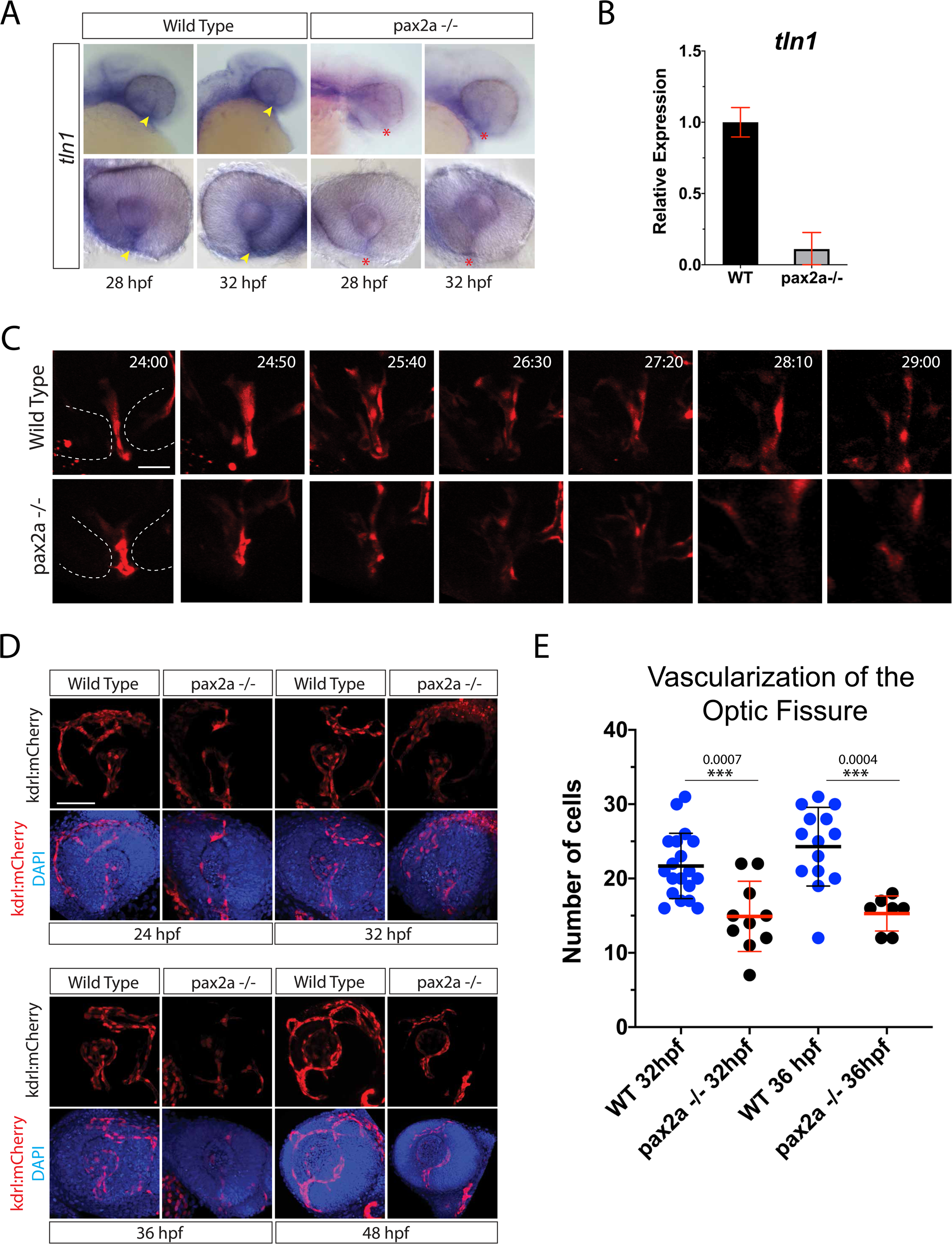
Pax2a is necessary for *tln1* expression and recruitment of vasculature into the optic fissure. **A)** Wholemount in situ hybridization comparing *tln1* expression between WT and pax2a^−/−^ embryos at 28 and 32hpf. *tln1* signal within the optic fissure (yellow arrowhead) appears reduced in Pax2a^−/−^ embryos (red *). **B)** qPCR analysis of *tln1* expression in WT and pax2a-/- embryos at 32hpf confirms a reduction in *tln1* expression. **C)** *In vivo* 4D confocal imaging of Tg[kdrl:mCherry] WT or pax2a^−/−^ embryos. Time lapse series depicting the region of the optic fissure (dotted white lines) and mCherry positive vasculature endothelial cells migrating through the fissure from 24-29hpf. Scale bar = 10um. **D)** Comparison of WT and pax2a^−/−^ vascularization during optic fissure fusion, 24-48hpf. 3D reconstructions of whole mount Tg[kdrl:mCherry] (red) WT or pax2a^−/−^ embryos with DAPI (blue) stained DNA. Scale bar = 50μm. **E)** Quantification of the number of mCherry positive cells from 3D confocal stacks within the region of the optic fissure. Individual embryo results are depicted. ANOVA p<0.0001.

### Inhibition of VEGF signaling impairs optic fissure fusion mechanics

Based on our discovery of impaired OF hyaloid vasculature in pax2a^−/−^ embryos, we next examined whether this phenomenon is associated with failure of optic fissure fusion. Hence, we turned our attention to vascular endothelial growth factor (VEGF) signaling. VEGF, the ligand for vascular endothelial growth factor receptor (VEGFR), is a prime candidate for regulating the migration and proliferation of hyaloid vasculature cells. In support of this notion, when examining expression patterns for VEGF ligands, vegfaa and ab, ba, c and d, we found *vegfaa, ab* and *c* to be expressed in the retina and periocular regions (Fig S4A). To inhibit VEGF signaling, we took advantage of the dorsomorphin derivative DMH4, which has been shown in zebrafish to selectively inhibit VEGF signaling independent from BMP (38). Based on published working concentrations, we conducted a dose response to examine DHM4 effects on hyaloid vasculature using Tg[kdrl:mCherry] as a readout (38). Treatment of embryos from 12-24hpf, ranging from 1-100μM, resulted in a dose dependent reduction of mCherry signal in the developing retina (Fig S4B). We decided to use the 100μM concentration for subsequent experiments as this concentration was able to completely inhibit vascularization of the retina up to 72hpf without any significant impact on overall embryo health (Fig 5A, S4C). Embryos were treated starting at 12hpf and examined for fissure fusion status via whole mount laminin IHC at 24, 32, 36, 48, 56 and 72hpf (Fig 5B). 3D confocal imaging revealed a persistence of laminin signal within the fissure up and including 72hpf in all embryos examined (Fig 5B). Measurement of the laminin signal generated very similar results to what we observed in pax2a^−/−^ embryos (Fig 5C). This is highlighted by retention of laminin signal at 48hpf and persistence of laminin in 100% (n=49) of embryos treated up to 72hpf (Fig 5B-C). In addition to laminin, we also imaged and quantified OF F-actin levels during the treatment (Fig 5B). Similar to pax2a^−/−^ embryos DMH4 treatment prevented the accumulation of OF F-actin between 24-32hpf (Fig 5D). This was in contrast to DMSO treated embryos which exhibited the accumulation of F-actin in the fissure between 24-32hpf like wildtype embryos. DMH4 treatment did not affect the size of the retina nor the apposition of retinal lobes (Fig S2). Lastly, we also observed a significant decrease of *tln1,* but not *pax2a,* expression in DMH4 treated embryos (Fig S4E,F). This suggests that *tln1* is likely expressed by the vasculature cells within the OF. Based on these findings, we conclude that inhibition of VEGF signaling results in failure of F-actin accumulation and basement membrane breakdown, presumably due to the absence of hayloid vasculature cells in the OF.

**Figure 5:**
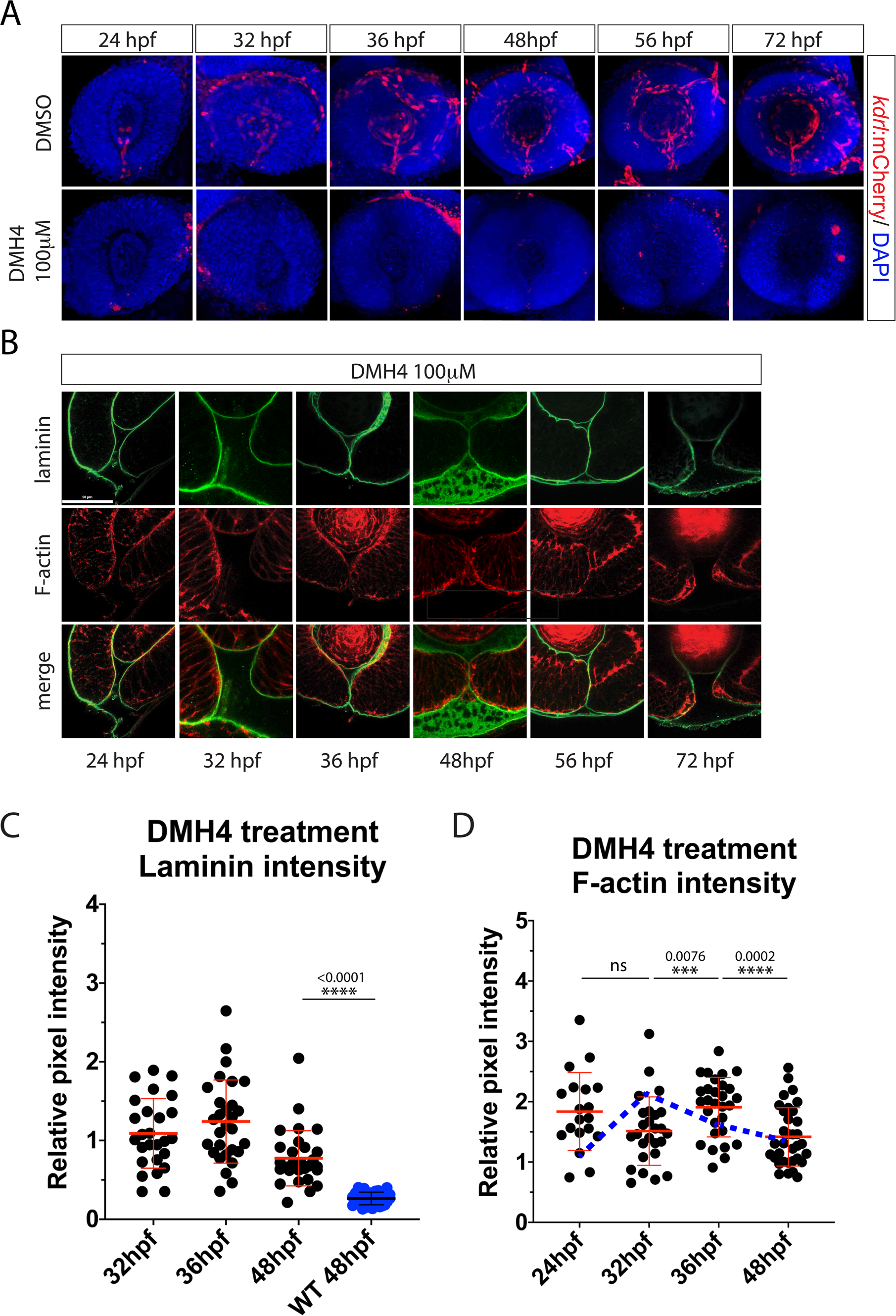
Inhibiting angiogenesis prevents optic fissure fusion. **A)** 3D confocal imaging of Tg[kdrl:mCherry] (red) embryos treated with DMSO or 100μM DMH4 between 24-72hpf. DNA was stained with DAPI (blue). Scale bar = 50μm. DMH4 treatment results in the absence of hyaloid vasculature the fissure and retina. **B)** Whole mount Immunohistochemistry was used to simultaneously visualize F-actin (red) and laminin (green) in DMH4 treated embryos from 24-72hpf. Central-proximal regions of the optic fissure are displayed. Treatment with DMH4 results in retention of laminin and optic fissure fusion failure. Scale bar = 50μm. **C)** Quantification of laminin signal intensity within the central-proximal region of the optic fissure, normalized to regions of laminin staining juxtaposed to the lens. Relative pixel intensities are displayed. ANOVA p<0.0001. **D)** Quantification of F-actin signal intensity (phalloidin staining) within the central-proximal region of the optic fissure, normalized to regions of F-actin signal within the lobe of the retina. Relative pixel intensities are displayed. A blue dotted line represents the WT trend in F-actin intensity. ANOVA p<0.0001.

### Anti-angiogenic protease *adamts1* is upregulated in OF of pax2a mutant embryos

Having established a connection between pax2a function and recruitment of hyaloid vasculature cells into the optic fissure, we next sought to examine the mechanism behind it. We therefore re-examined our transcriptomic comparison of WT and pax2a^−/−^ eyes, with a focus on angiogenic regulation. This new examination revealed misexpression of *a disintegrin and metalloproteinase with thrombospondin motifs 1* (adamts1), a secreted protease known to be involved in regulating VEGF signaling (39). Being a member of the ADAMTS family, adamts1 contains three thrombospondin motifs enabling it to directly bind and sequester VEGF (31). Furthermore, the protease activity of adamts1 has bene shown to target thrombospondin 1 and 2 for cleavage, ultimately liberating their active forms which bind to and block VEGF from binding the VEGF receptor (30). Our RNAseq data indicated that *adamts1* expression was upregulated almost two-fold in pax2a^−/−^ eyes (table 1). This would suggest that the expression of *adamts1*, and therefore its anti-angiogenic activity, is normally kept in check by pax2a. Pax2a has not been reported to harbor transcriptionally repressive function, so this may involve an indirect mechanism. To confirm our RNAseq results we performed WISH, examining *adamts1* expression at 28 and 36hpf. In agreement with our transcriptomic analysis, we observed an upregulation in *adamts1* in the ventral region of pax2a^−/−^ eyes, specifically surrounding the optic fissure (Fig 6A). A statistically significant increase in *adamts1* expression was also confirmed using qPCR (Fig 6B). To test whether upregulation of *adamts1* expression would have direct effects on hyaloid vascularization and optic fissure fusion we injected Tg[*kdrl*:mCherry] embryos with *adamts1* mRNA. We subsequently examined hyaloid vasculature using 3D confocal imaging between 24-48hpf. Injection of *adamts1* mRNA resulted in a significant decrease in the number of *kdrl*:mCherry cells found within the optic fissure (Fig 6B, C, Movie 5). The observed decrease mirrored the results from pax2a^−/−^ embryos (Fig 6C). Our findings therefore suggest that *adamts1* expression needs to be tightly regulated in order to ensure proper vascularization of the retina, including the OF. Furthermore, the observed up-regulation of *adamts1* expression in pax2a^−/−^ embryos may explain the resulting OF hyaloid vasculature deficiency.

**Figure 6:**
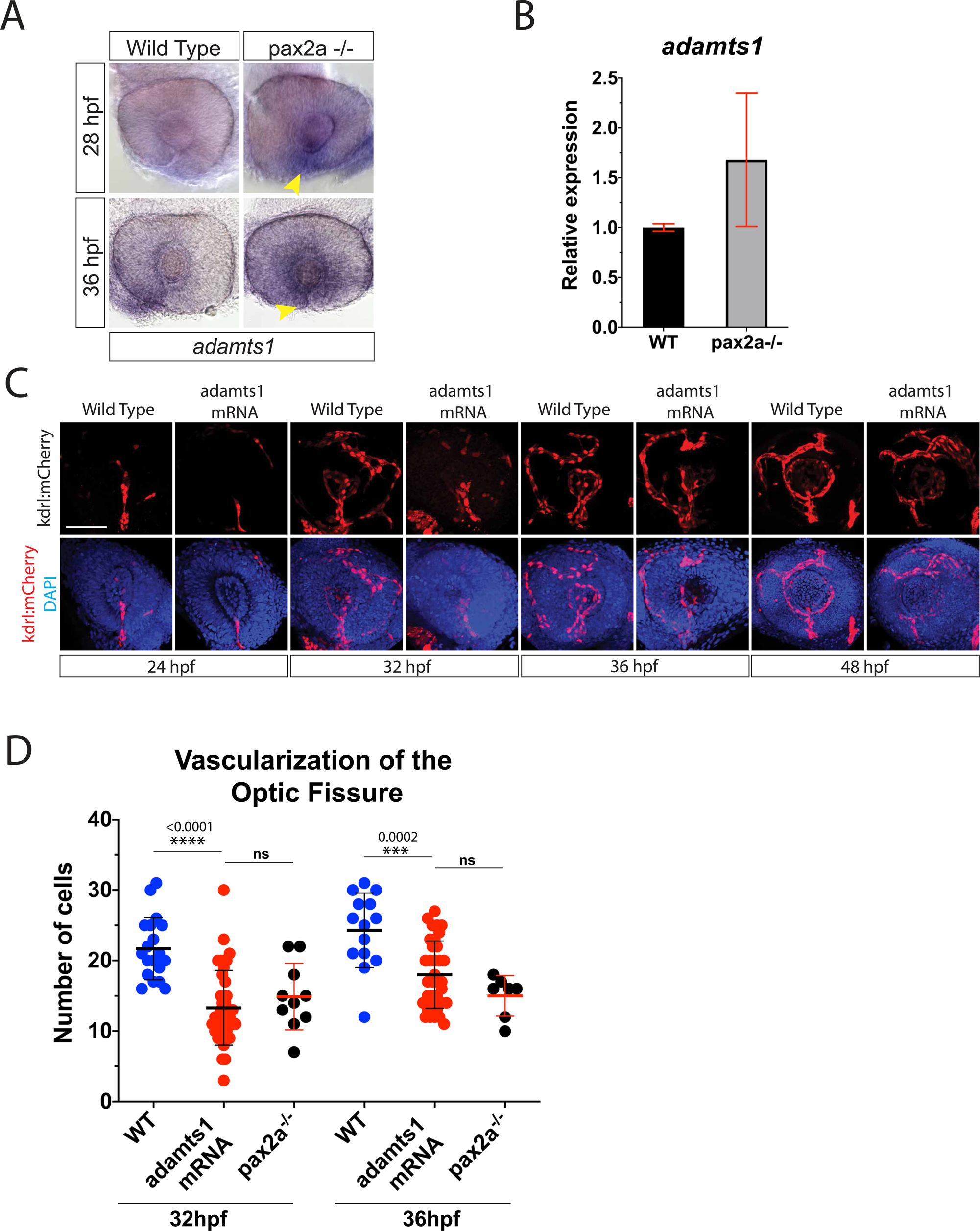
Adamts1 inhibits retinal vascularization. **A)** Wholemount in situ hybridization comparing *adamts1* expression between WT and pax2a^−/−^ embryos at 28 and 32hpf. *adamts1* signal within the optic fissure (yellow arrowhead) is appears increased in Pax2a^−/−^ embryos (red *). **B)** qPCR analysis confirms an increase in *adamts1* expression in pax2a-/- embryos. **C)** Comparison of WT and adamts1 mRNA injected embryo vascularization during optic fissure fusion, 24-48hpf. 3D reconstructions of whole mount Tg[kdrl:mCherry] (red) WT or adamts1 mRNA injected embryos with DNA stained by DAPI (blue). Scale bar = 50μm. **D)** Quantification of mCherry positive cells from 3D confocal stacks within the region of the optic fissure. Individual embryo results are depicted. * p<0.001, ANOVA p<0.0001.

### Overexpression of *adamts1* inhibits optic fissure fusion mechanics

Having mimicked the vasculature phenotype of pax2a mutants by over expressing *adamts1*, we next determined whether it had any effects on OF fusion. To do so, we performed whole mount laminin and F-actin staining in *adamts1* mRNA injected embryos at 24, 32, 36, and 48hpf (Fig 7A). Confocal imaging of the fissure indicated that *adamts1* mRNA injection also phenocopied pax2a^−/−^ associated persistence of laminin and therefore failure of optic fissure fusion. Quantified levels of laminin persisting at 48hpf was similar to pax2a^−/−^ embryos (Fig 7B). Similar to pax2a^−/−^ and DMH4 treated embryos, we again noted a lack F-actin accumulation between 24-32hpf in *adamts1* overexpressing embryos (Fig 7C). As was observed with pax2a^−/−^, timing for apposition of the lobes was not affected in adamts1 overexpressing embryos (Fig S2). Taken together, our data suggest that maintaining proper levels of *adamts1* expression is necessary to enable VEGF signaling and subsequent recruitment of hyaloid vasculature required for initiation of OF BM remodeling.

**Figure 7:**
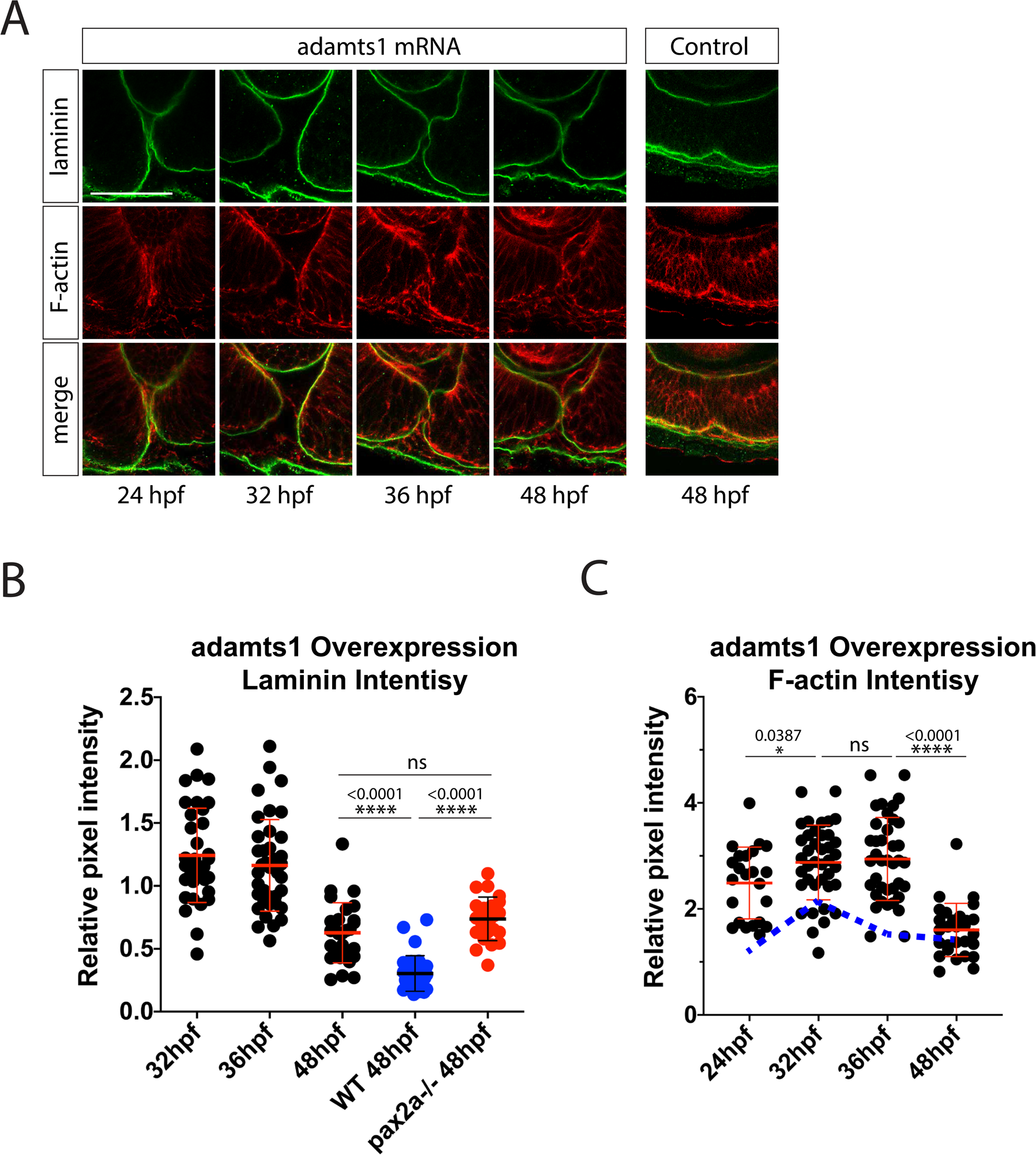
*Adamts1* misregulation leads to failure of optic fissure fusion. **A)** Whole mount Immunohistochemistry was used to simultaneously visualize F-actin (red) and laminin (green) during optic fissure fusion, 24-72hpf in adamts1 mRNA injected embryos. Central-proximal regions of the optic fissure are displayed. Scale bar = 50μm. **B)** Quantification of laminin signal intensity within the central-proximal regions of the optic fissure, normalized to regions of laminin staining juxtaposed to the lens. Relative pixel intensities are displayed. ANOVA p<0.0001. **C)** Quantification of F-actin signal intensity (phalloidin staining) within the central-proximal regions of the optic fissure, normalized to regions of F-actin signal within the lobe of the retina. Relative pixel intensities are displayed. A blue dotted line represents the WT trend in F-actin intensity over time. ANOVA p<0.0001.

### Hyaloid vasculature is the source of mmp2 necessary for OF BM remodeling

The above data confirm and support a model where hyaloid vasculature drives or initiates the OF fusion process. However, a missing key to this model is the mechanism by which vasculature cells induce fissure fusion. James *et al*, along with others, have suggested vasculature cells may be a source of BM remodeling enzymes, such as matrix metaloproteases (mmp). To investigate this further, we used WISH to examine mmp expression within the OF between 24-48hpf. Our data indicated that *mmp2, 14a* and *14b* were expressed within the fissure between 28-36hpf (Fig S5 and Famulski lab unpublished data). Mmp2 had recently been associated with OF fusion in the mouse while evidence from an mmp2 metalloproteinase activity probe in zebrafish indicated mmp2 activity is present in the developing eye and likely OF (40, 41). mmp14 is an activator of mmp2 and its co-expression with mmp2 within the fissure suggests mmp2 is in fact active (42, 43). To test whether mmp2 fits within our model we assayed *mmp2, mmp14a and mmp14b* expression in pax2a^−/−^ and DMH4 treated embryos. In both cases, OF expression of *mmp2, mmp14a and mmp14b* was reduced as observed by WISH and confirmed by qPCR (Fig 8A, B). This suggested that all three are expressed by the hyaloid vasculature cells. To confirm this, we performed 2 color WISH for *mmp2* and *kdrl*. Confocal imaging verified that *mmp2* expression is co-localized with that of *kdrl* within the OF (Fig 8C). Conversely, when performing 2 color WISH for *mmp2* and *rorB*, a retina specific probe, we did not detect co-localization of the signals. We therefore conclude that the source of mmp2, 14a and 14b enzymes in the OF are hyaloid vasculature cells. Finally, to determine whether mmp2 activity is necessary for OF fusion we treated embryos with ARP101, an mmp2 specific inhibitor previously shown to be effective in zebrafish (44). Treating embryos from 24-48hpf inhibited OF BM remodeling in a dose dependent manner (Fig 8D,E, S6). When compared to DMSO, embryos treated with 20μM ARP101 either completely or partially retain their OF BM up to 48hpf. Mmp2 activity is therefore necessary for BM remodeling in the OF. To determine when mmp2 activity is required we began the ARP101 treatment between 24-32hpf and assayed BM status at 48hpf (Fig 8F). Treatments started later than 30hpf had little to no effect on OF BM remodeling, indicating that mmp2 activity is required between ∼24-30hpf (Fig 8F). This finding also correlates with the observed timing of *mmp2* expression in the OF (Fig S5). However, because we cannot determine exactly how quickly ARP101 inhibits enzymatic activity in our assay, active mmp2 may persist in the OF longer than 30hpf.

**Figure 8:**
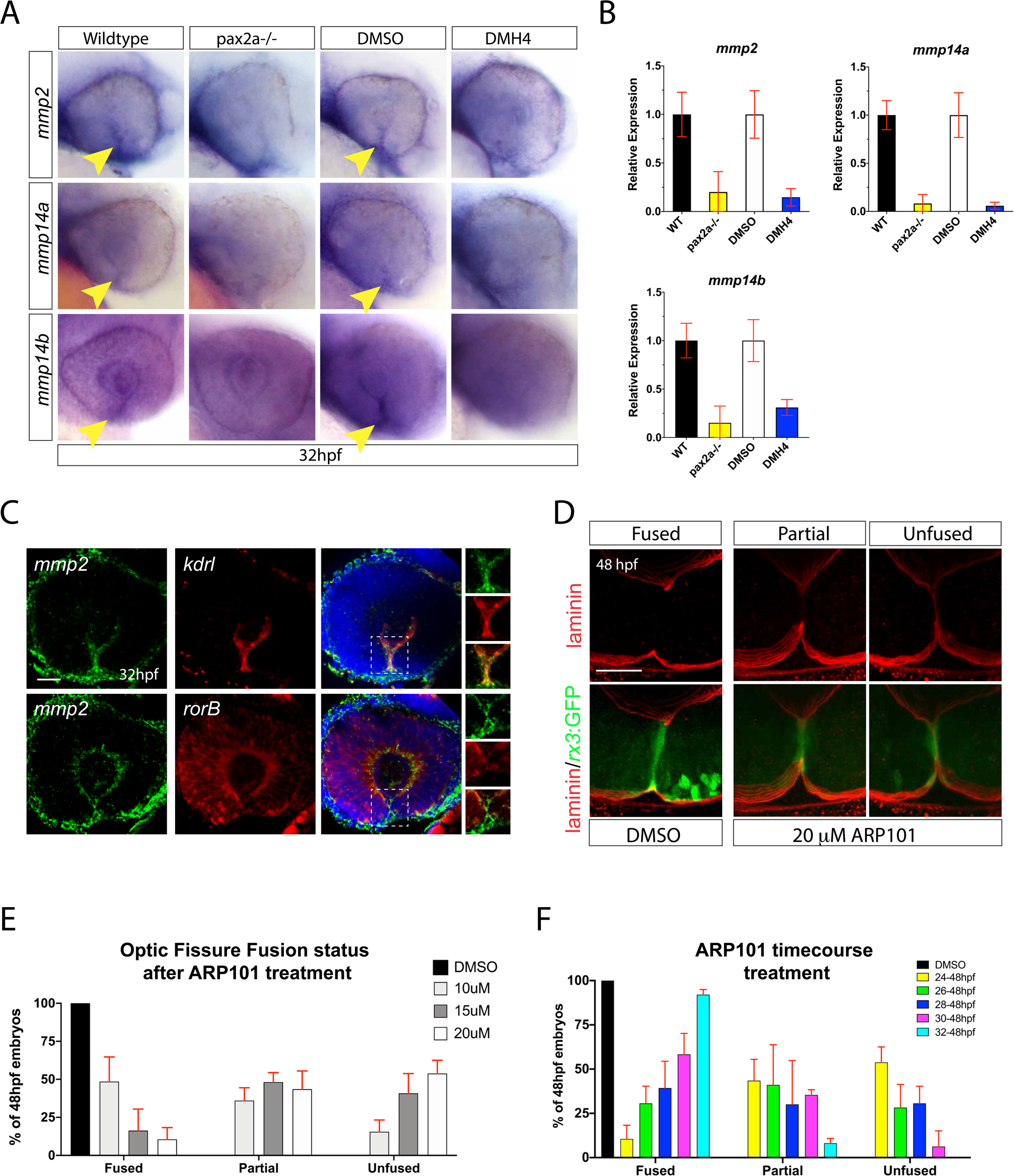
Hyaloid vasculature is the source of mmp2 activity required for optic fissure BM remodeling. **A)** Wholemount in situ hybridization comparing *mmp2, mmp14a* and *mmp14b* expression in WT vs pax2a^−/−^ and DMSO vs DMH4 treated embryos at 32hpf. *Mmp2, 14a and 14b* signal within the optic fissure (yellow arrowhead) appears decreased in pax2a^−/−^ and DMH4 treated embryos. **B)** qPCR analysis confirms a decrease in expression of *mmp2, 14a* and *14b* in both pax2a^−/−^ and DMH4 treated embryos. **C)** Two color wholemount in situ hybridization simultaneously examining *mmp2* and *kdrl*, or *mmp2* and *rorB* expression at 32hpf. DNA was stained with DAPI. Scale bar = 50μm. Clear overlap of signal is observed for *mmp2* and *kdrl*, but not *mmp2* and *rorB*. **D)** Whole mount Immunohistochemistry was used to visualize laminin (red) in ARP101 treated Tg[*rx3*:GFP] embryos at 48hpf. Central-proximal regions of the optic fissure are displayed. **E)** Quantification of 48hpf ARP101 treated embryos for fissure fusion (absence of laminin signal), partial fusion (partial retention of laminin) or failure to fuse (retention of laminin throughout the fissure). n= 41 (10mM), 38 (15mM), 42 (20mM) ANOVA p<0.0001. **F)** Quantification of 48hpf ARP101 treated embryos for fissure fusion, partial fusion or failure to fuse after various treatment initiation times from 24-32hpf. n= 42 (24-48hpf), 29 (26-48hpf), 29 (28-48hpf), 23 (30-48hpf), 37 (32-48hpf). ANOVA p<0.0001.

## DISCUSSION

Studies of optic fissure fusion dating back several decades have been suggesting a direct connection between the fusion process and hyaloid vasculature found within the fissure. In fact, this hypothesis has been recently strengthened by data showing that a reduction of hyaloid vasculature in the fissure, or removal of optic vesicles from sources of vasculature inhibits or significantly delays fusion (22, 28). In our study, we have characterized a pax2a driven mechanism that ensures proper vascularization of the OF by restricting anti-angiogenic activity of adamts1 to enable hyaloid vasculature invasion of the OF and delivery of BM remodelers mmp2, 14a and 14b. In conclusion, our findings further validate the notion that hyaloid vasculature is an active component of the machinery driving OF fusion.

Several recent reports, including this one, have comprehensively characterized optic fissure fusion timing (21–24). Work from zebrafish, mice and chick all point to an orderly progression involving: **1)** retinal growth and cellular rearrangement leading to nasal and temporal retinal lobe apposition, **2)** invasion of the fissure by endothelial and neural crest cells forming the hyaloid vasculature system, **3)** cellular signaling, either between retinal rim cells or between rim cells and the migrating vasculature cells, **4)** remodeling or removal of the basement membrane to enable physical connection of the rim cells and subsequent formation of a continuous retinal epithelial sheet via re-polarization and cell-cell adhesion. Step 1 has been nicely characterized in a few recent publications outlining the flow of retinal cells and morphological formation of the fissure (45–47). For step 2, several reports have carefully characterized the formation of the hyaloid vasculature system, including migration of hyaloid vasculature precursor cells into the fissure as soon as it forms (1–3). Importantly, perturbation of this process, or removal of the developing eye from its source, has been shown to lead to fissure fusion failure (22, 28). To date, steps 3 and 4 are the least understood. Recent work from our lab has characterized the composition of core BM components within the fissure (23). Additionally, work from other labs has identified several molecular components associated with cell-cell adhesion and epithelial sheet fusion to function within the fissure, including *α*-catenin, n-cadherin, and netrin (22, 24, 48–51). However, the timing and the molecular mechanism organizing and regulating these components remains uncharacterized. Finally, the elephant in the room has always been the identity of the BM remodeling mechanism. To date, only adamts16 has been functionally examined in context of fissure fusion, mmp2 activity was detected during mouse OF fusion and mmp23bb was implicated in the fusion process from transcriptomic data (41, 50, 52).

While attempting to address the mechanistic aspects of steps 3 and 4 using the pax2a^noi^ model we first uncovered a relationship between F-actin and BM remodeling. Work by *James et al. 2016* suggested that F-actin accumulation may be indicative of hyaloid cell interaction with retinal rim cells. During our detailed time course analysis, we discovered that BM remodeling is in fact preceded by an increase in F-actin signal within the OF (Fig 1). When assayed in pax2a^−/−^ embryos the F-actin accumulation is absent while the hyaloid vasculature in the OF is also diminished. The timing of F-actin accumulation coincides with the active migration of hyaloid vasculature cells through the fissure (Fig 4). Furthermore, we show pharmacologically, via VEGF inhibition, that hyaloid vasculature is necessary for the accumulation of F-actin and OF BM remodeling. Transcriptionally, we show that the decrease in hayloid vasculature in pax2a^−/−^ embryos coincides with a significant decrease in *tln1* expression. Tln1 is a key regulator of endothelial cell migration, recently shown to be directly involved in OF fusion (22). In fact, loss of tln1 function has been previously shown to decrease in fissure associated vasculature and inhibit OF BM remodeling (22). We also show a significant decrease in *tln1* expression upon VEGF inhibition (DMH4 treatment), suggesting that *tln1* expression within the OF is associated with hayloid vasculature. Talin1 is known to be a direct link between the actin cytoskeleton and the BM via integrin (53, 54), and has been associated with the formation of adhesion junctions (55, 56). One could therefore hypothesize several different models for its role in optic fissure fusion. We predict that tln1 is regulating the ability of hayloid vasculature cells to migrate properly through the OF, however we cannot rule out the possibility that tln1 directly participates in the OF fusion process. The absence of tln1 expression is most likely a correlation to the decrease of hayloid vasculature cells within the fissure of pax2a^−/−^ embryos. Our transcriptomic analysis was performed using whole eye tissue, which would have included the hayloid vasculature. As such, the decrease in *tln1* expression is likely indicative of decreased numbers of hayloid vasculature cells which require *tln1* expression for proper migration into the fissure and or proliferation.

Our study strongly supports the notion that vasculature plays an integral part in OF fusion. As such, modulation of angiogenesis represents a potential mechanism to ensure proper vascularization of the OF. In our transcriptomic comparison of WT and pax2a mutants we did in fact discover a connection to angiogenesis. In the absence of pax2a function we observed an upregulation of *adamts1*. Examining *adamts1* expression in the eye further confirmed an upregulation of expression in the ventral regions pax2a^−/−^ retinas. Encoding 3 thrombospondin (thsb) motifs able to directly bind and sequester VEGF, in addition to targeting thsb1 and 2 for cleavage into their active VEGF inhibiting form, adamts1 is well known for it’s anti-angiogenic function in cancer, aortic and renal biology (57). In our study, we show that upregulating adamts1 expression reduces the number of hyaloid vasculature cells in the OF. Up regulation of adamts1 also prevents the accumulation of F-actin and subsequent BM remodeling. Taken together we propose that the reduction in hayloid vasculature observed in pax2a^−/−^ embryos likely stems from an upregulation of adamts1 leading to suppressed VEGF signaling and therefore limited vascularization of the OF.

The last aspect of our proposed model pertains to the mechanism of OF BM remodeling. It had been suggested over the years that vasculature may be the source of BM remodeling activity during OF fusion. To that end, we have discovered that *mmp2*, *mmp14a* and *mmp14b* are all expressed within the hayloid vasculature during OF fusion. In fact, timing of their expression matches precisely with the initiation of BM remodeling. When we examine pax2a mutants or inhibit hayloid vasculature completely we no longer detect expression of either of these mmps. Furthermore, using ARP101, a specific mmp2 inhibitor, we show that mmp2 activity is necessary for OF BM remodeling and this activity is specifically required between 26-32hpf. Mmp2 has been implicated in OF fusion previously. Mouse studies have shown mmp2 expression within macrophages residing in the OF. Furthermore, recent examination of mmp2 activity, using a reporter construct, indicates that mmp2 is active in the eye and likely the OF (40). Mmp14 is a known to activate mmp2 activity by cleaving the inhibitory pro-peptide of mmp2. As such, co-expression of mmp2 and 14 is further indicative of mmp2 playing a functional role in OF BM remodeling.

To date, the only BM remodeling enzyme to be associated with OF fusion is adamts16. Loss of adamts16 function in zebrafish, via morpholino, led to a coloboma like phenotype which the authors credited to the inability to degrade laminin in the fissure (52). We have not been able to reproduce adamts16 expression within the fissure. Recent studies in mice, zebrafish and chick have compared expression in OF cells pre, during and post fusion (24, 50, 58, 59). Surprisingly, none of these studies identified any obvious candidates for carrying out BM remodeling, including mmp2 and mmp14. However, it remains possible that the OF tissue examined lacked the hayloid vasculature and therefore prevented the identification of mmp2 and mmp14 as candidates. Lastly, expression of *mmp2, 14a* and *14b* is transient in the fissure which may have contributed to their OF expression being missed by other groups. Our data indicate mmp2 activity is necessary for BM remodeling of the OF, but does not rule out a role for additional proteases. There may be several proteases involved, such as adamts16, mmp2 and others yet inhibition of just one could trigger failure of the remodeling process and lead to fissure fusion failure. Continuing studies into matrix protease activity will be needed to fully characterize the remodeling mechanisms during OF fusion.

In conclusion, we present a new model for the mechanism of OF BM remodeling where pax2a restricts expression of *adamts1* in the ventral retina to enable hayloid vasculature invasion of the OF. Once in the fissure, vasculature cells express *mmp2, 14a* and *14b* to initiate BM remodeling. As soon as fusion begins, vasculature becomes restricted from the fissure and ultimately the two retinal lobes fuse. Our model aligns with the molecular events observed in the fissure, including accumulation of F-actin at the time of vasculature migration through the fissure (24-28hpf), expression of mmps (26-36hpf) and subsequent BM remodeling (32-48hpf). For future studies it will be important to understand how pax2a regulates *adamts1* expression and to examine whether mutations in *adamts1*, *mmp2* or *mmp14* are potential markers for coloboma.

## Supplemental Material

**Figure S1:**
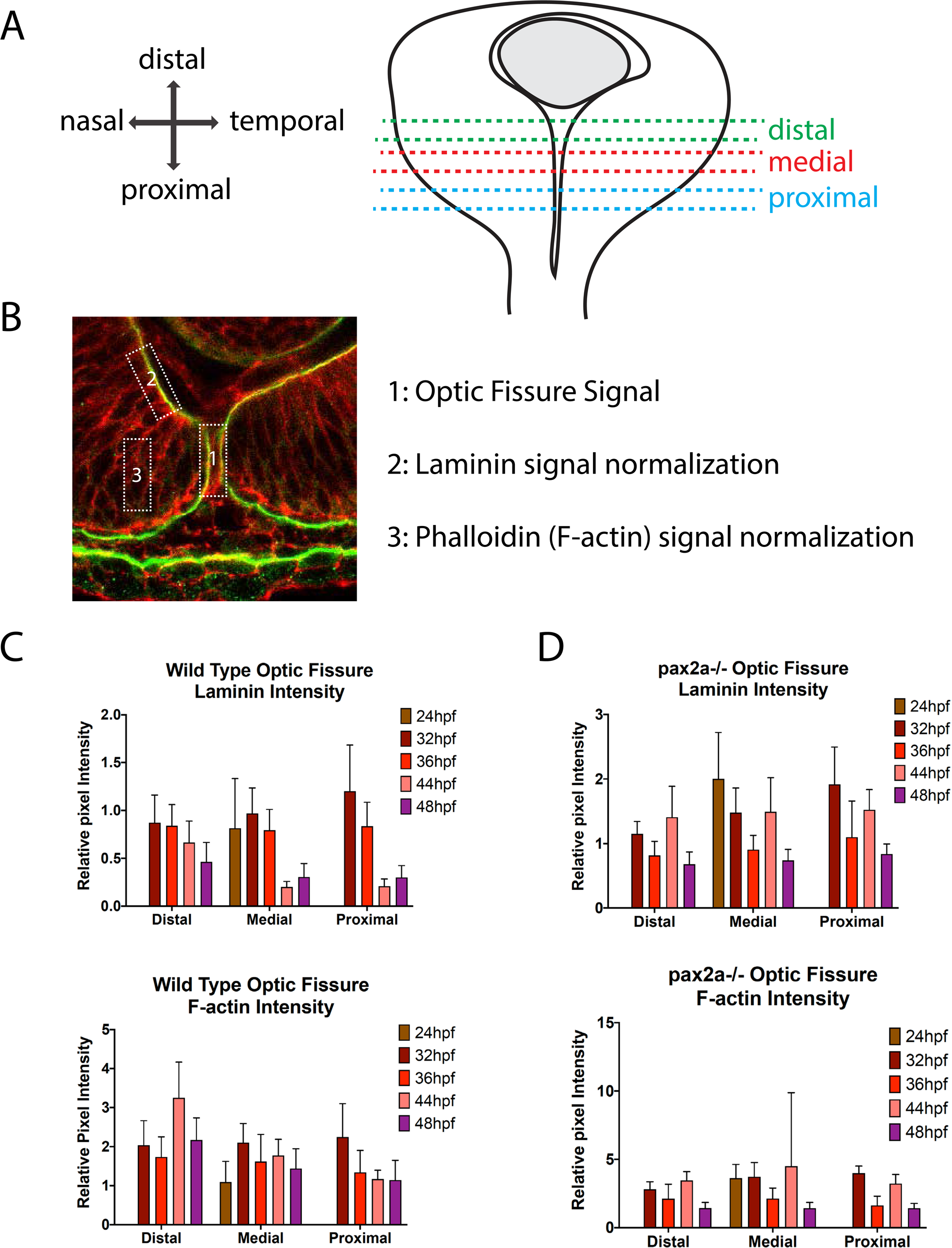
Optic Fissure quantification. **A)** Graphical representation of the region of OF analyzed using confocal microscopy. **B**) Sample image depicting regions of the OF (1), laminin normalization (2) and F-actin normalization (3) used for signal intensity quantification**. C)** Quantification of laminin and F-actin signal intensity in wild type embryos within the distal, medial and proximal regions of the optic fissure, normalized to regions of laminin staining juxtaposed to the lens and F-actin signal within the lobe of the retina. Relative pixel intensities are displayed. ANOVA p<0.0001. **D)** Quantification of laminin and F-actin signal intensity in pax2a^−/−^ embryos within the distal, medial and proximal regions of the optic fissure, normalized to regions of laminin staining juxtaposed to the lens and F-actin signal within the lobe of the retina. Relative pixel intensities are displayed. ANOVA p<0.0001.

**Figure S2:**
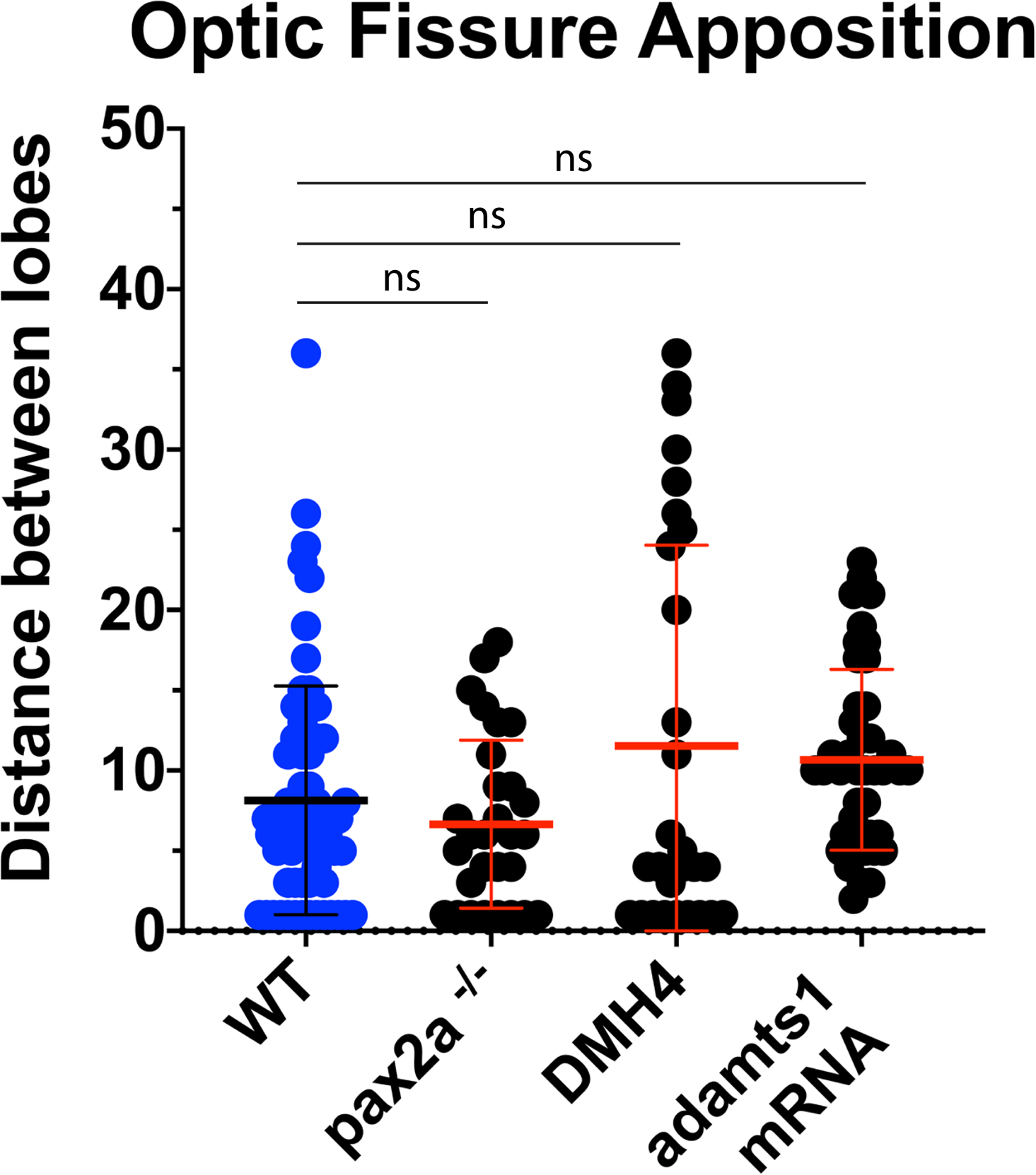
Optic Fissure apposition measurements. Measurements of the distance between retinal lobes (apposition) in WT, pax2a^−/−^, DMH4 treated or ADAMTS1 mRNA injected embryos at 48hpf. Measurements were made using laminin staining as reference for edges of retinal lobes. Distance was measured as pixels.

**Figure S3:**
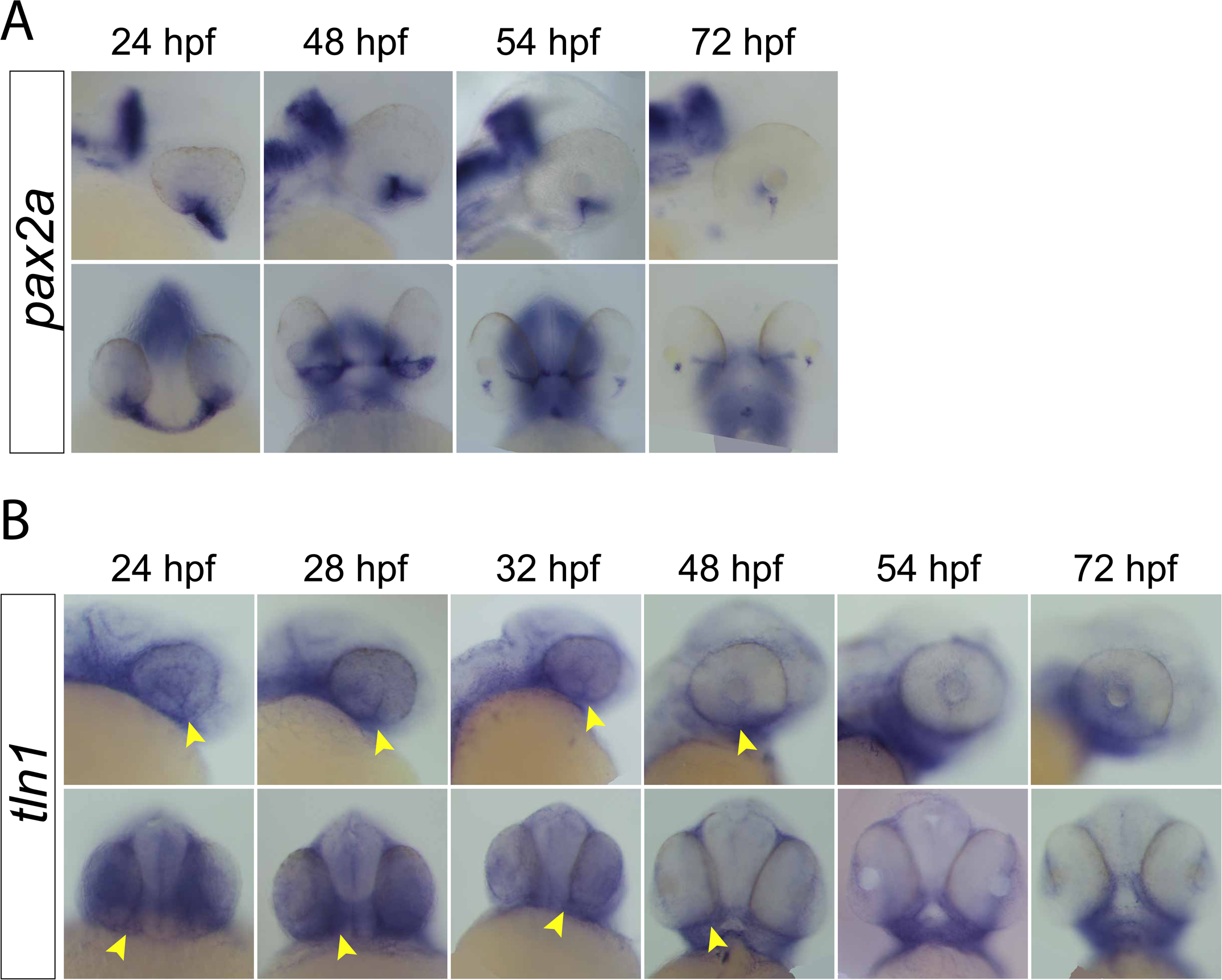
*pax2a* and *tln1* expression during development. **A)** Whole mount in situ hybridization of *pax2a* probe at 24, 48, 54 and 72hpf. Lateral (top) and ventral (bottom) images depicting optic fissure expression are shown. Pax2a expression persists in the fissure up to 54hpf. **B**) Whole mount in situ hybridization of *tln1* probe at 24, 48, 54 and 72hpf. Lateral (top) and ventral (bottom) images depicting optic fissure expression are shown. *tln1* expression is detected in the optic fissure from 24-48hpf (yellow arrowheads).

**Figure S4:**
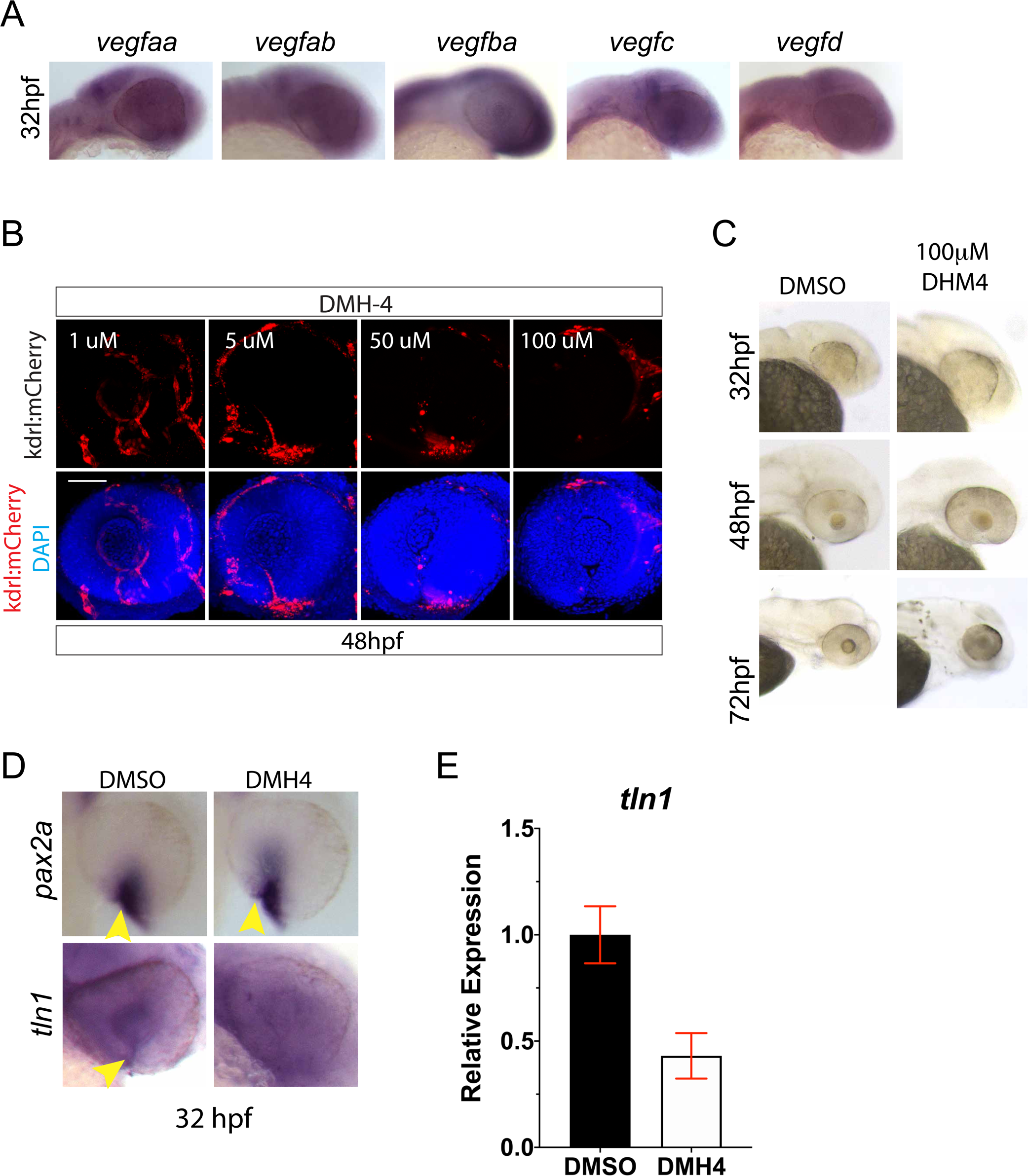
DMH4 dose response. **A)** Whole mount in situ hybridization of *vegfaa, ab, ba, c* and *d* probes at 32hpf. **B)** 3D reconstructions of 48hpf whole mount Tg[kdrl:mCherry] (red) embryos treated with 1, 5, 50 or 100uM DMH4. DNA stained with DAPI (blue). Increasing concentration of DMH4 eliminates mCherry expressing cells from the optic fissure and retina. Scale bar = 50μm. **C)** Brightfield images of DMSO or DMH4 treated embryos at 24, 48 and 72hpf. **D)** Whole mount in situ hybridization of *pax2a* or *tln1* probe at 32hpf in DMSO or DMH4 treated embryos. OF expression is indicated with a yellow arrowhead. DMH4 treatment does not appear to alter *pax2a* expression but eliminates *tln1* expression from the OF. **E)** qPCR results for *tln1* expression in DMH4 treated embryos.

**Figure S5:**
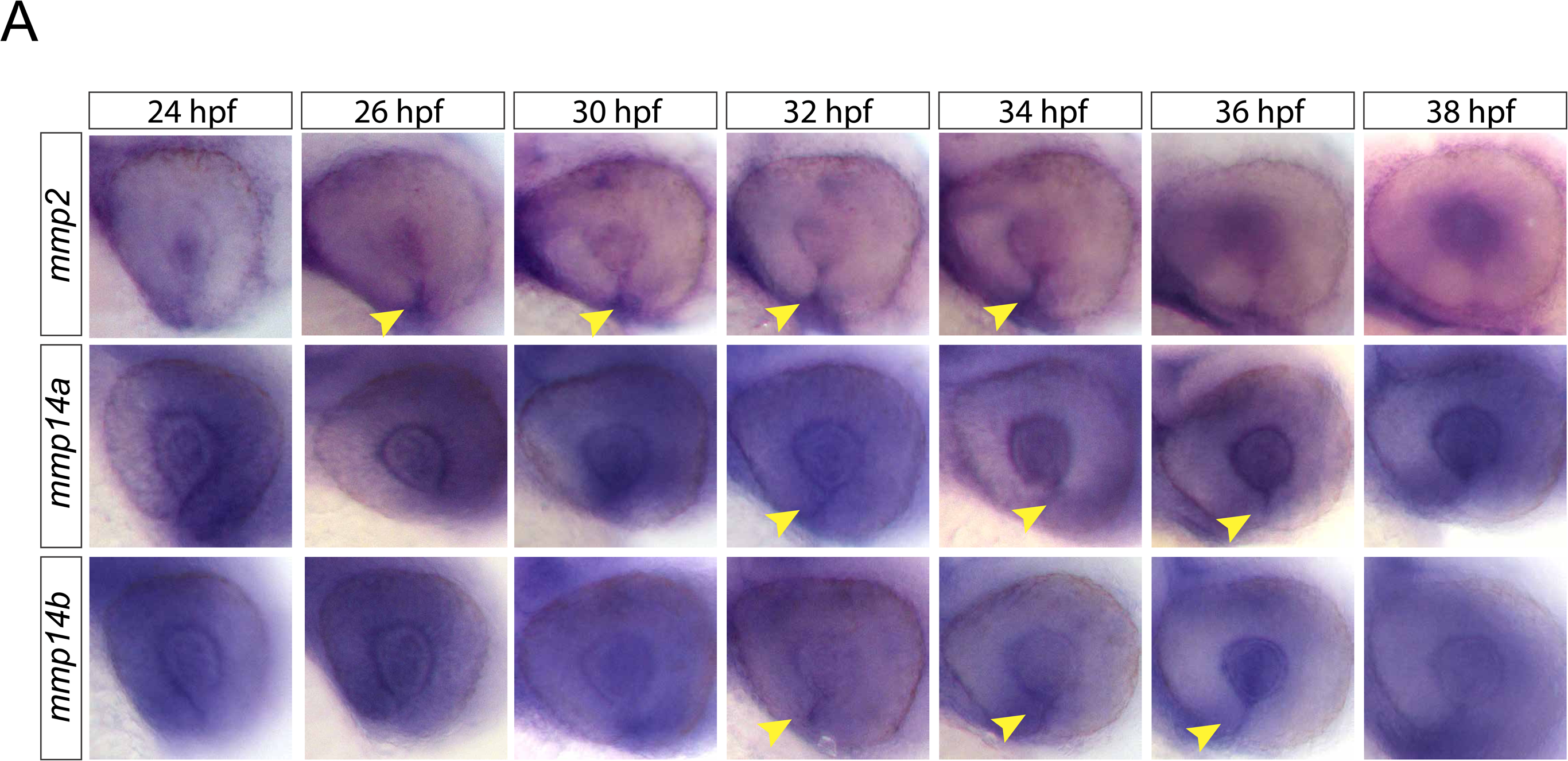
Time course of *mmp2, mmp14a* and *mmp14b* ocular expression. Whole mount in situ hybridization of *mmp2, mmp14a, mmp14b* probes at 24, 26, 30 32, 34, 36 and 38hpf. *Mmp2* expression is present in the optic fissure from 26-34hpf (yellow arrowheads). *Mmp14a* expression is present in the optic fissure from 32-36hpf(yellow arrowheads). *Mmp14b* expression is present in the optic fissure from 32-36hpf (yellow arrowheads).

**Figure S6:**
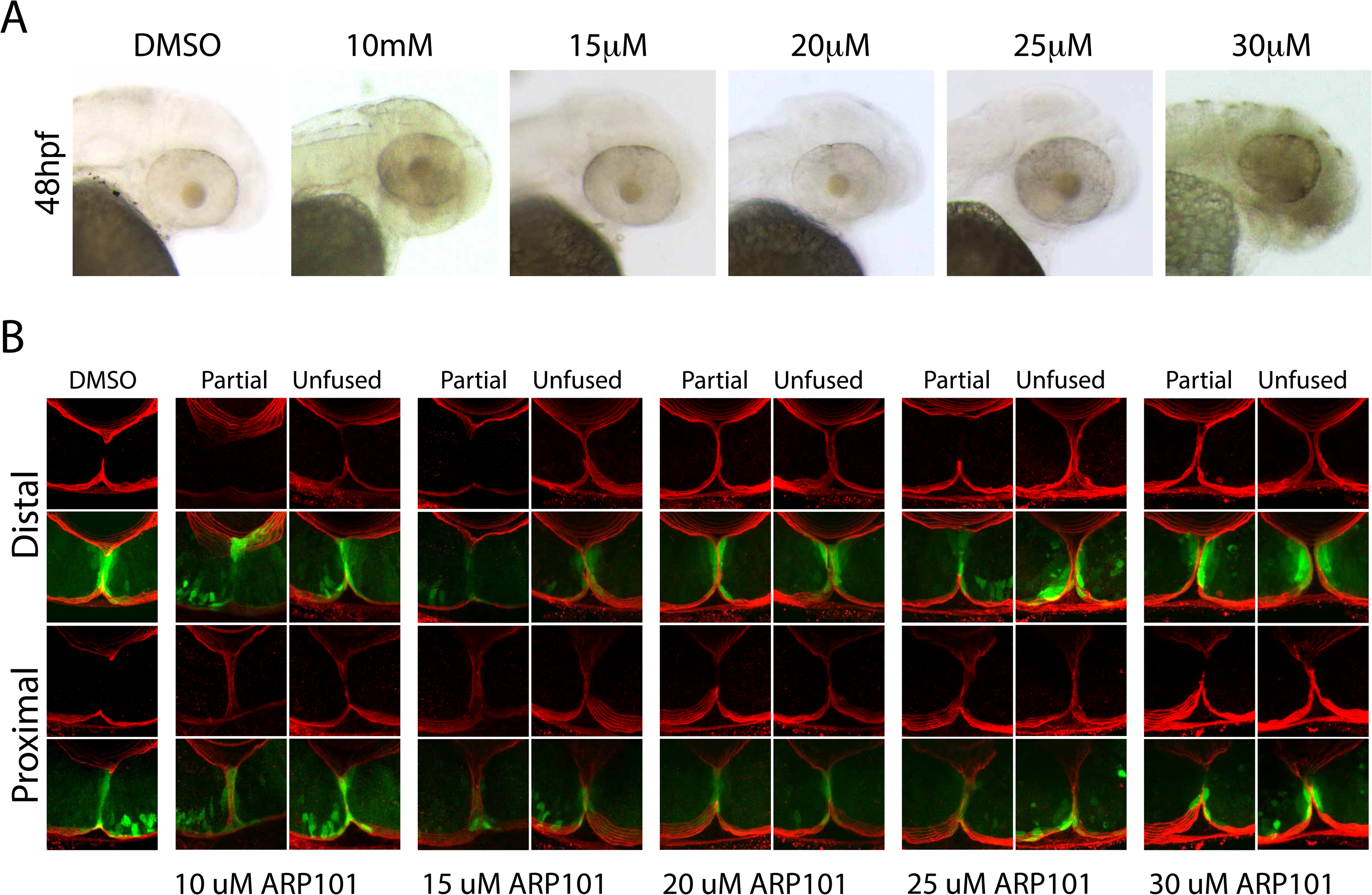
ARP101 treatment dose response. **A)** Brightfield images of DMSO and ARP treated embryos at 48hpf. Concentrations of ARP101 exceeding 20uM result in toxic effects. **B)** Whole mount Immunohistochemistry was used to visualize laminin (red) in ARP101 treated Tg[*rx3*:GFP] embryos at 48hpf. Central-proximal and distal regions of the optic fissure are displayed.

Movie 1: WT Tg[kdrl:mCherry] migration within the optic fissure, 24-30hpf

Movie 2: pax2a^−/−^ Tg[kdrl:mCherry] migration within the optic fissure, 24-30hpf

Movie 3: 3D rotation of WT Tg[kdrl:mCherry] embryo at 48hpf

Movie 4: 3D rotation of pax2a^−/−^ Tg[kdrl:mCherry] embryo at 48hpf

Movie 5: 3D rotation of 48hpf Tg[kdrl:mCherry] embryo injected with ADAMTS1 mRNA

Movie 6: 3D rotation of 48hpf Tg[kdrl:mCherry] embryo injected with ADAMTS1 morpholino

Table 1: 95^th^ percentile RNA sequencing comparison

Table 2: Raw data RNA sequencing comparison

Table 3: Primer sequences

## MATERIALS AND METHODS

### Zebrafish and embryo maintenance

Zebrafish were maintained using husbandry procedures approved by University of Kentucky IACUC committee. Embryos were kept at 28.5°C in E3 embryo media. AB and TL strains were used as wild-type, Tg[*kdrl*:mCherry] transgenic line was used to visualize retinal vascularization (60), Tg[*rx3*:GFP] was used to visualize retinal cells (61).

Pax2^noi^ embryos were a gift from Dr. Gregory-Evans. Genotyping analysis was conducted by amplifying the region of gDNA with the noi mutation using the forward primer: 5’- CTCGCTCTGCCTCCATGATTG3-’and the reverse: 5’- GGCACTGAAAGAGCACAGG -3’. The resultant 460bp amplicon was digested with TaqI (NEB) which would recognize and digest the WT allele sequence but not the mutant allele.

### Immunohistochemistry (IHC)

Dechorionated embryos were fixed with 4% PFA in PBS at room temperature for 3h and washed with PBST 4 times for 5 minutes. Embryos were then permeabilized with Proteinase K, 30μg/mL 10 minutes for 24-28hpf, 50μg/mL 15-20 minutes for 32-48hpf and 75μg/mL 20 minutes for 56-72hpf, washed 2 times in PBST for 5 minutes and blocked overnight at 4°C with 10% sheep serum, 0.8% Triton X-100 and 1% BSA in PBS. Primary mouse anti-laminin antibody (ThermoFisher – 1:100) in blocking buffer (1% sheep serum, 1% BSA and 0.8% Triton X-100 in PBS) were incubated overnight at 4°C and washed 5 times in PBST for 15 minutes. Secondary antibody, goat anti-rabbit (Alexa Fluor® 488 – Abcam – 1:1000), DAPI 1:1000, and phalloidin (Alexa Fluor® 555 - 1:50) were incubated overnight at 4°C in the dark. Tg[*kdrl*:mCherry] embryos were treated with proteinase K as described above and stained with DAPI 1:1000 overnight. The embryos were washed 2 times in PBST for 15 minutes and visualized using a Nikon C2+ confocal microscope equipped with a 40X (1.15NA) water immersion objective. Embryos were embedded in 1.2% low melting point agarose on glass bottom 35mm dishes (Fluorodish, World Precision Instruments). Images were captured in steps of 3.5 microns for a total of 31.5 microns using Nikon Elements software. Image adjustment, such as cropping and brightness/contrast was performed using Adobe Photoshop.

### Analysis of Fluorescence signal

Fiji software (https://fiji.sc) was used to measure the fluorescence intensity of laminin and phalloidin signal from raw image data. In order to account for variability in staining, normalization values were measured for laminin and actin pixel intensity where an area directly outside of the choroid fissure was measured and a ratio was generated between the two values (Fig S1B). In cases where the fissure edges were farther apart then the size of the box used for analysis, the box was divided into two and used to measure fluorescence intensity on each side of the fissure. Measured values from both boxes was added and then normalized. For Tg[krdl:mCherry], 3D reconstructions of the optic fissure were generated and individual cells were counted (from the opening of the OF through the back of the lens).

### Stats

Student’s t-test was used to compare individual time points. One-way ANOVA was used to analyze across treatments. Graphs are displayed as mean +/- standard deviation. Analysis was performed using Prism8 graphing software (GraphPad).

### Total RNA Sequencing

WT and Pax2^−/−^ embryos were dissected and collected at 48hpf and RNA was extracted from dissected eyes using trizol. Pax2a^−/−^ embryos were phenotyped by distinct heart defects only observed in pax2a^−/−^ embryos, as verified by previous genotyping experiments. Dissected eyes were suspended in 1 mL of trizol and sheared with a 22-gauge needle. Samples were incubated at room temperature for 5 minutes, then 200 *μ*L of chloroform was added and vortexed for 1 minute. Samples were then centrifuged at 12,000 g for 15 minutes at 4℃. Aqueous phase was removed and put into fresh RNAse free tube. 3 *μ*L of Glycoblue was added to samples and vortexed for 5 seconds. 500 *μ*L of 100% Isopropanol was then added and vortexed for an additional 10 seconds. Samples then were incubated at room temperature for 10 minutes. Subsequently, samples were centrifuged at 12,000 g for 10 minutes at 4℃. Supernatant was removed and 1 mL of 75% EtOH was added and vortexed for 10 seconds. Samples were then centrifuged at 7,500 g for 5 minutes at 4℃. Supernatant was removed and samples then underwent a pulse centrifugation and any remaining supernatant was removed. Samples were air-dried for 7 minutes at room temperature under a fume hood. 20 *μ*L of RNAse free ddH_2_O was added to the samples and mixed until pellet dissolved. Finally, samples were incubated at 60℃ for 12 minutes and stored at -80℃.

The RNA then underwent a DNase treatment using the DNA-free Kit. 2 *μ*L of 10X DNaseI Buffer, and 1 *μ*L of rDNaseI was added to the entire 20 *μ*L sample. Samples were incubated at 37℃ for 20 minutes, and then 2 *μ*L of DNase Inactivation Reagent was added. Samples were then incubated at room temperature for 2 minutes, mixing samples 3 times. Samples were centrifuged at 10,000 g for 1.5 minutes. Supernatant was then removed and placed into fresh RNase free tube and stored at -80℃.

Purified total RNA was sent to Applied Biological Materials Inc. for Illumina sequencing (PE150bp, 30-40 million reads/sample). Bioinformatic analyses were completed using RSEM. Those transcripts which had a greater than 0.95 posterior probability of being differentially expressed were considered significant. Panther (http://pantherdb.org) was used for gene ontology and pie chart generation.

### Whole-mount in situ hybridization (WISH)

Whole-mount in situ hybridization was performed as previously described (62). RNA probes were generated using PCR with T7 promoter sequence linkers and subsequently transcribed [DIG or FITC labeled] using T7 polymerase (Roche). Primer sequences are all found in Table 3. Images were captured using a Nikon Digital sight DS-Fi2 camera mounted on a Nikon SZM800 stereo scope using Elements software. Dissected eyes from 24, to 72hpf embryos were mounted in 70% glycerol and imaged under DIC using a Nikon TiE compound microscope equipped with a 20X (0.7NA) objective and Elements software. Image adjustment, such as cropping and brightness/contrast was performed using Adobe Photoshop.

### qPCR analysis

32hpf embryos were anesthetized with tricane, tail tips were collected for genotyping and the heads, dissected just posterior of the eyes, were fixed in RNAlater. After genotyping, embryos corresponding to WT and pax2a-/- were pooled, 5-10 embryos, and total RNA isolated using a RNAaqueous kit (Ambion). DMSO and DMH4 treated embryos were harvested in the same fashion absent any genotyping. qPCR was performed as previously described (63).

### 2 color fluorescent in situ hybridization

Fluorescent whole mount in situ hybridization was performed as previously described (64). Images were collected using a Nikon C2+ confocal microscope with a 20X (0.95NA) objective and images were adjusted for brightness and contrast using Adobe Photoshop.

### Live imaging analysis

Live imaging of Tg[kdrl:mCherry] embryos was conducted using a Nikon C2+ confocal microscope equipped with at 20X (0.95NA) water immersion objective. 22hpf embryos were imbedded in 1.1% low gelling agarose in 1-inch glass bottomed Flourodish cell culture dishes (World Precision Instruments) and covered in embryo media, 3-amino benzoic acidethylester (tricaine) to anaesthetize the embryos and 1-phenyl 2-thiourea (PTU) to inhibit pigmentation. Z-stacks 75μm thick with a step size of 2.5μm were captured over the course of 6 hours at 10 minute intervals. The time lapse data were reconstructed in 3D using Elements software. Image adjustment, such as cropping and brightness/contrast was performed using Adobe Photoshop. After imaging, embryos were removed and genotyped.

### Inhibitor treatments

Embryos were incubated in embryo media with 5, 25, 50, or 100 *μ*M of DMH-4 (Sigma Life Science) in DMSO starting at 12hpf. Fresh DHM4 containing media was added every 12hpf for timepoints past 24hpf. For ARP101 (Tocris) treatments embryos were dechorionated and treated with 10, 15, 20, 25 or 30μM ARP101 in fish water from 24 to 48hpf, unless stated otherwise.

### Cloning and mRNA synthesis

Full coding domain sequences for *adamts1* (NCBI reference sequence XM_021475923.1), were amplified from 24hpf zebrafish cDNA using primers 5’- CTCGAGTCAACAGGGAGTCAGATTGC-3’ for *adamts1*, cloned into pGEMT (Promega), digested with AsiSI/XhoI, BamHI/XhoI respectively and subsequently cloned into pCS2+. All constructs were verified by sanger sequencing (Eurofinsgenomics). mRNA was synthesized from linearized pCS2 constructs using SP6 mMessage mMachine kit (Ambion) and purified using YM-50 Microcon columns (Amicon, Millipore).

### Ethics Statement

The use of zebrafish in this study was approved by the University of Kentucky IACUC committee, Institutional PHS Assurance #D16-00217 (A3336-01) with a protocol number: 2015-1370.

## ACKNOWLEDGEMENTS

This work for supported by faculty start-up funds provided to JKF from the department of Biology, University of Kentucky. MLW was supported by the Lyman T Johnson scholarship from the University of Kentucky. WPP was supported by Brazilian National Council for Scientific and Technological Development (CNPq) under grant number 202970/2014-0. We thank Dr. Jeramiah Smith for assistance with bioinformatic analysis of our RNAseq data and members of the Famulski lab for helpful discussions.

